# Trait empathy modulates music-related functional connectivity

**DOI:** 10.1101/2021.08.18.456484

**Authors:** Vishnu Moorthigari, Emily Carlson, Petri Toiviainen, Peter Vuust, Elvira Brattico, Vinoo Alluri

## Abstract

It has been well established through behavioural studies that empathy significantly influences music perception. Such individual differences typically manifest as variability in whole brain functional connectivity patterns. To date, nobody has examined the modulatory effect of empathy on functional connectivity patterns during continuous music listening. In the present study, we seek to investigate the global and local connectivity patterns of 36 participants whose fMRI scanning was done by employing the naturalistic paradigm wherein they listened to a continuous piece of music. We used graph-based measures of functional connectivity to identify how cognitive and affective components of empathy modulate functional connectivity. Partial correlation between Eigenvector centrality and measures of empathy showed that cognitive empathy is associated with higher centrality in the sensorimotor regions responsible for motor mimicry while affective empathy showed higher centrality in regions related to auditory affect processing. We furthermore identified a left-hemispheric dominance in the modulatory effect of affective empathy particularly in the Orbitofrontal cortex and the temporal pole. Results are discussed in relation to various theoretical models of empathy and music cognition.

## Introduction

Music is a modality of human interaction which appears in all known cultures, serving primarily social, relational and emotional functions (1). Music allows us to express and convey emotions and to share intentions and experiences that are aesthetic, embodied and participatory in nature (2–6). Indeed, the power of music to induce emotions has been explained with the human capacity to understand and share the emotional states of others; that is, to feel empathy for others (7, 8). This capacity to feel empathy is fundamental for social functioning, including prosocial behaviour (9–11), and its dysfunction is implicated in many different psychological and behavioural disorders, such as Autism Spectrum Disorders (12, 13), schizophrenic disorders, and Borderline Personality Disorder (14–16). Investigation of the relationship between music and empathy has suggested that individual differences in the tendency to empathize relate to susceptibility to emotion induction by music (17), to enjoyment of sad music (18), to a tendency to move spontaneously when hearing music (19), and to a tendency to interact with others while moving to music (20).

Hence, the ability to feel others’ emotions and to understand their intentions is important to music interaction, and might be one of the mechanisms behind music’s ability to elicit emotions in listeners and performers alike. Indeed, Omigie (8) points to empathy, along with expectancy, as a core mechanism allowing for emotional responses to music receiving most consensus by psychology and neuroscience scholars. Scherer and Zentner (21) suggested that emotional responses to music rely on the listener’s identification with the composer or performer and an empathic reaction to their expressed emotion, while Watts (22) suggests that, in the absence of an observed performer, music can represent a virtual persona with whom listeners may empathize. Leman (2), with a perspective rooted in embodied cognition, has suggested empathy in music listening involves the listeners’ covert (mental) or overt (bodily) imitation of ‘moving sonic forms.’ A similar idea comes from Juslin and Vastfjall (4), under the label ‘emotional contagion,’ which they suggest entails the listener internally mimicking the music’s affect, which leads to the experience of a similar emotion as that expressed by the music. In a similar vein, Overy (23) introduced the Shared Affective Motion Experience (SAME) model, in which music is conceptualized as an “intentional, hierarchically organized sequence of expressive motor acts,” emphasizing the social interactive nature of music making. This model suggests that the activation of the same or similar neural networks in an agent and observer, that is, a performer and listener, allows empathy to be a route to emotional response to music (24).

Although no single accepted theoretical model of empathy currently exists, it is commonly conceptualized as including affective and cognitive components as well as a mechanism for tracking self-feelings and other-feelings (25–27). Accordingly, various researchers (28, 29) have chosen to study empathy as consisting of two distinct but interrelated systems – Cognitive and Affective empathy. Cognitive empathy is associated closely with Theory of Mind and describes an individual’s ability to infer another’s mental state including aspects such as their knowledge or beliefs, whereas Affective empathy is associated with understanding another’s affective state, sometimes related to state-matching, emotional contagion or experiences of empathic concern (26, 30). Over and covert motor mimicry are believed to be among the key mechanisms supporting empathic abilities. Individuals differ in their tendency to empathise in certain situations and in certain ways; such ‘trait empathy’ is traditionally measured using self-report scales. Measures differ in the degree to which they focus on and differentiate between various aspects of empathy; the Balanced Emotional Empathy Scale (BEES) (31), for example, focuses on emotional empathy, while the Empathy Quotient (13) integrates cognitive and affective aspects of empathy, and the Interpersonal Reactivity Index measures cognitive and affective aspects of empathy using four subscales: Fantasy Seeking (FS), Perspective Taking (PT), Empathic concern (EC) and Personal Distress (PD) (32).

This distinction between cognitive and emotional aspects of empathy has not been largely examined in the context of music. Hargreaves and Colman (1981) proposed two possible music listening strategies used by listeners - ‘Objective-analytic’ and ‘Affective,’ which is suggestive of the Cognitive-Affective distinction in empathy (33). Recent evidence additionally suggests two motivational dimensions underlying aesthetic appreciation of music and elicited pleasure, that is, relaxational-sensational and social-contemplative (34) which might be central in modulating individual responses. The relaxational-sensational dimension can be described as an affective, potentially low-level physiological response, whereas the social-contemplative dimension describes processing music as more of a cerebral endeavour, which involves processing the music in a more analytical fashion. This distinction is interesting in the light of Affective vs Cognitive empathy and suggests that an individual automatically processes music employing one of the two approaches, which may be modulated by his or her empathic predispositions.

Over the last couple of decades, multiple studies have examined the neural substrates empathy primarily using visual stimuli. One such study by Chakrabarti et al. (35) found that while participants viewed facial images depicting various emotions their Empathy Quotient (EQ) was correlated positively with activity in the precuneus, prefrontal cortex, inferior frontal gyrus (IFG) and premotor cortex, and negatively with activity in the insula and temporal lobe. In another study conducted by Nummenmaa et al. (36) the authors categorised the visual stimuli presented to participants into Cognitive and Emotional empathy blocks and asked the participants to empathize with people in the scene. Their results showed that stimuli depicting Emotional empathy resulted in greater activity in the Thalamus, Fusiform gyrus, inferior parietal lobule, middle occipital gyrus and parahippocampal gyrus whereas scenes depicting Cognitive empathy resulted in activation in the medial prefrontal cortex. Furthermore, their study did not find any differential activation in the premotor cortex, which the authors attributed to the stimuli being static goal-directed actions resulting in both types of stimuli having similar activations in the region. Fan et al. (2011) performed a meta-analysis including forty fMRI studies related to empathy, a majority of which focussed on pain-related visual stimuli such as evaluation of emotional faces, or imagining oneself in specific emotional situations. They identified the so-called Core Empathy Network (CEmN) (37) which includes median and anterior cingulate cortex (MCC, ACC), insula extending to the IFG, supplementary motor area (SMA), premotor cortex, orbital frontal cortex (OFC), thalamus, midbrain and the precentral gyrus. Following this, they further divided the tasks into those that involved Affective-perceptual empathy (which is analogous to Affective empathy) and those that involved Cognitive-evaluative empathy (analogous to Cognitive empathy). In addition to the above common regions of activation, they found that Affective empathy involved greater activation in the midbrain, right Anterior Insula, Right Dorsal-medial Thalamus (DMT) and the right ACC. On the other hand, Cognitive-evaluative empathy showed greater activation for the left OFC, left MCC and left DMT. However, all the studies included in this meta-analysis were solely activation-based utilising a segregated approach, wherein brain regions or voxels are examined independently. In light of recent studies such as that of Gratton et al. (38) showing that individual-specific factors dominate the variability of whole brain network topology, it has become important to augment the segregated approach with an integrative approach wherein global and local connectivity patterns are also examined.

Thus far, only two studies (39, 40) have tried to identify the neural mechanisms underlying the processing of music that are associated with empathy, with the goal of testing the core hypothesis of whether emotions felt in response to music are derived from empathy predispositions. Wallmark et al. employed a more controlled paradigm that required participants to passively listen to short musical segments (2s/16s). In their study, the FS and PT subscales of the IRI were considered to make up cognitive empathy while EC and PD sub-scales made up affective empathy. Their results with very short stimuli (2s) showed that greater cognitive empathy was associated with increased activity in the SMA, somatosensory cortex as well as the ACC and for the 16s stimuli, they observed increased activation in parts of the prefrontal cortex and tempo-parietal junction (TPJ). For Affective empathy, their results showed increased activations in the IFG, OFC, superior temporal gyrus (STG), Inferior parietal lobule (IPL), somatosensory cortex, cerebellum and parts of the midbrain. In the second study, Sachs et al. utilised the naturalistic paradigm, wherein the participants were exposed to stimuli in a setting closer to everyday experiences, such as listening to music without performing any other task, allowing for more natural responses which cannot be evoked in a lab setting (41, 42). However, their study was specific to sad music and focused primarily only on the Fantasy Seeking dimension of cognitive empathy, leaving it open whether other dimensions of empathy would also modulate brain responses to music. In the first part of their study, their results showed increased ISC in clusters in the bilateral auditory cortices extending to the insula, ACC and the IFG for all participants while they listened to a full length music piece. In addition to the above, their study involved dividing the participants into two groups (high and low empathy) based on participants’ Fantasy seeking scores and subsequently evaluating the differences in Inter-subject correlation (ISC) of the two groups. For the second part of their study, they showed that participants scoring high on FS had significantly greater ISC in the left auditory cortex, dorsal medial prefrontal cortex (DMPFC), precuneus and parts of the visual cortex whereas participants scoring low on FS showed significantly greater ISC in the insula and the caudate. It was proposed that this finding is relevant in the context of participants becoming cognitively engaged with the music as it unfolded. Although both these studies shed some light on the associations between empathy and neural processing of music, they have focused solely on activation-based patterns. No studies till date have looked at how empathic traits manifests as functional connectivity, which is representative of brain states, during music listening. The current study attempts to fill this gap by examining how functional connectivity is related to individual differences in empathy during music listening. To this end, we utilize graph-theoretical approaches to model whole-brain connectivity. In recent years, computational advances in graph theory have provided a strong foundation to study patterns of functional connectivity in naturalistic fMRI data (43, 44), making it a powerful tool to understand global functional network organisation. In the context of continuous music listening, graph-based approaches such as node degree centrality have previously been used to identify central hubs and how their organization is modulated by participants’ musical expertise (45). Another study by Toiviainen et al. (46) used dynamic graph based measures such as eigenvector centrality to understand how musical beat salience affects the organisation of functional networks.

In the current study, we first divided the participants into two groups based on their empathy score, by performing a median-split, to check for the presence of group-level differences in ISC. For this first part of the study, in line with previous studies (39, 47) we hypothesize that at the whole-group level, significant mean ISC would primarily be centered around the bilateral auditory cortices. Second, for the ISC difference between high and low empathy groups, we hypothesize that participants belonging to high Cognitive empathy groups (Fantasy Seeking in particular) would have greater ISC scores in the temporal cortex, precuneus and the visual cortex as observed in Sachs et al. Thus far, there haven’t been any studies examining ISC differences based on participants’ affective empathy in the context of music listening. However, based on studies using non-musical stimuli, we hypothesize that the group scoring high on Affective empathy subscales (i.e., EC and to a lesser extent PD) would show greater ISC in the thalamus and the insula.

Next, we employ a model-free approach of capturing functional connectivity via graph-theory. We identified central hubs related to empathy which are key in organizing global connectivity during naturalistic music listening. Due to the exploratory nature of our connectivity analyses we do not have any pre-defined hypotheses. Nevertheless, at a global-level, we expect that for individuals scoring high on the cognitive empathy scales, regions belonging to the premotor cortex (SMA and pre-/post-central gyri) as well as the ACC and insula would play a central role. Whereas for those scoring high on affective empathy scales, we hypothesize that regions belonging to the corticolimbic system such as the temporal pole and IFG involved in the encoding of musical-affect related would play a central role. Finally, we examine empathy-driven differences in local connectivity through modularity. Since the modularity analysis was performed based on the regions identified to be central in global connectivity, we cannot have a priori hypotheses.

## Methods

### fMRI data acquisition

The study is part of the broad “Tun-teet” research protocol involving multi-dimensional brain measures, psychological tests and behavioural data on audition, emotion and musical behaviour. The protocol aimed at studying the neural sources of individual differences in sound-induced emotions. All experimental procedures for this protocol were approved by the Coordinating Ethics Committee of the Hospital District of Helsinki and Uusimaa (approval number: 315/13/03/00/11, obtained on March the 11th, 2012). Furthermore, all procedures were conducted in agreement with the ethical principles of the Declaration of Helsinki. Further details on this protocol can be found from (46, 48). The same fMRI dataset, having the same fMRI scanning and preprocessing pipeline but tested for other hypotheses and devoid of the questionnaires considered here, has been published in (45, 46, 49). The current study is a re-analysis of the dataset in combination with previously-unpublished behavioural questionnaire data on the same sample.

#### Participants

Thirty-six healthy participants (age 28.6 *±* 8.9, 20 females) with no history of neurological or psychological disorders participated in the fMRI experiment. They were recruited among university students or working professionals (firm employees or entrepreneurs). Before being admitted to the research, the participants were screened for inclusion criteria before admission to the experiment (no ferromagnetic material in their body; no tattoo or recent permanent coloring; no pregnancy or breastfeeding; no chronic pharmacological medication; no claustrophobia). The participants had variable levels of music education, with a median of 3.5 years and an interquartile range of 11.25 years of formal music education.

#### Stimulus & task

Participants’ fMRI brain responses were acquired while they listened to an Argentine tango by Astor Piazzolla, Adiós Nonino (8-min long). This stimulus was chosen because it includes notable acoustic variation and expresses several different emotions, some sections being more melancholic, while others are energetic and exciting. Participants’ task was to attentively listen to the music delivered via high-quality MR-compatible insert earphones (Avotec, Stuart, FL, USA) while keeping their eyes open. To encourage participants to actively attend to the music, before stimuli were presented, we informed the participants that they would have to answer several questions after each stimulus by talking on the intercom. These questions concerned affective ratings of the stimuli on a 5-point Likert scale (among which liking and familiarity will be considered for this study), and were recorded with paper and pencil by the experimenter and later checked for consistency with other affective ratings taken in a separate session. These behavioural measures, as well as the video monitoring of participants concord in showing that participants were able to maintain attention during music listening. The sound level of the stimuli was individually adjusted so that they were audible above the scanner noise but the volume stayed within safety limits (below 80 dB).

#### Scanning

Scanning was performed using a 3T MAGNETOM Skyra whole-body scanner (Siemens Healthcare, Erlangen, Germany) and a standard 32-channel head-neck coil, at the Advanced Magnetic Imaging (AMI) Centre (Aalto University, Espoo, Finland). Using a single-shot gradient echo planar imaging (EPI) sequence thirty-three oblique slices (field of view: 192×192 mm; 64×64 matrix; slice thickness: 4 mm, interslice skip: 0 mm; echo time: 32 ms; flip angle: 75°; voxel size: 2×2×2 mm^3^) were acquired every 2s, providing whole-brain coverage per participant. T1-weighted structural images (176 slices; field of view: 256×256 mm; matrix: 256×256; slice thickness: 1 mm; interslice skip: 0 mm; pulse sequence: MPRAGE) were also collected for individual coregistration.

#### Preprocessing

Functional MRI scans were preprocessed on a Matlab platform using SPM8 (Statistical Parametric Mapping), VBM5 for SPM (Voxel Based Morphometry; Wellcome Department of Imaging Neuroscience, London, UK), and customized scripts developed by the present authors. For each participant, low-resolution images were realigned on six dimensions using rigid body transformations (translation and rotation corrections did not exceed 2 mm and 2°, respectively), segmented into grey matter, white matter, and cerebrospinal fluid, and registered to the corresponding segmented high-resolution T1-weighted structural images. These were in turn normalized to the MNI (Montreal Neurological Institute) segmented standard a priori tissue templates using a 12-parameter affine transformation. Functional images were then blurred to best accommodate anatomical and functional variations across participants as well as to enhance the signal-to-noise by means of spatial smoothing using an 8 mm full-width-at-half-maximum Gaussian filter. Movement-related variance components in fMRI time series resulting from residual motion artifacts, assessed by the six parameters of the rigid body transformation in the realignment stage were regressed out from each voxel time series. Following this, spline interpolation was used to detrend the fMRI data. Next, temporal filtering was performed by Gaussian smoothing (kernel width: 4 s), as it provides a good compromise between efficiency and bias (50).

### Behavioural Measures

Among the questionnaires filled in at Biomag laboratory, we included the Interpersonal Reactivity Index (IRI) (51), which gauges trait empathy scores. Unlike several other scales, the IRI includes four subscales which measure different components of empathy, in line with Decety and Jackson’s (2004) model of empathy as involving multiple, dissociable systems (52). The IRI consists of 28 items, answered using a five-point Likert scale. The sub-scales of the IRI consist of Perspective-taking (PT), Fantasy-Seeking (FS), Empathic Concern (EC), and Personal Distress (PD), each of which is made up of seven items. These four subscales have also been analysed under a two-dimensional model comprising Emotional (EC, PD) and Cognitive (FS, PT) empathy (40, 53, 54), this is in line with the view that empathy consists of two distinct subsystems - the affective system which involuntarily processes the emotions of another person, and the cognitive system which voluntarily simulates the emotions felt by another (54). However, a confirmatory factor analysis (55) showed that the two-factor split does not represent the underlying structure of the IRI, suggesting these factors represent a more complex underlying structure than given by a two-factor solution. Nevertheless, these four subscales can and have been conceptually grouped in previous studies (56); PT and FS are similar in that PT involves consciously taking the perspective of another person, while FS involves taking the perspective of an imaginary other person, as one might do in reading fiction. On the affective side, EC describes the tendency to feel sympathy or concern for others, while PD involves the tendency to experience negative emotions in response to another’s suffering. The IRI has been used in a wide range of research, for example relating IRI subscales to differences in brain structure (57) and demonstrating similar relationships between personality and trait empathy across diverse cultures (58). The IRI is used in the current study due to its usage and validation in multiple neuroscience studies (59–61).

### Inter-subject correlation

We used whole-brain inter-subject correlation (ISC), proposed by Hasson et al. (Hasson2004) to perform a preliminary analysis of the fMRI data. This was done by computing the voxel-wise Pearson’s correlation between the fMRI time series of every pair of participants, yielding an ISC map for every binary participant combination (for a total of 630 ISC maps). To verify the consistency of the participants’ fMRI responses, a group-level mean-ISC map was created by averaging the resultant ISC maps.

Following this, we divided the participants into low and high empathy groups by performing a median-split for each of the four IRI subscales. Mean ISC was then computed separately for each group using the aforementioned procedure. Thereafter, the voxel-wise ISC contrasts between the high and low empathy groups for each subscale were calculated by subtracting the mean ISC of the low empathy group from the high empathy group. Statistical significance of the resultant ISC contrast maps were calculated from a null-distribution of voxel-wise ISC difference values generated by performing 2000 iterations of bootstrapping with replacement from the original ISC maps (62). Refer to Figure 1 for an overview.

**Fig. 1.**
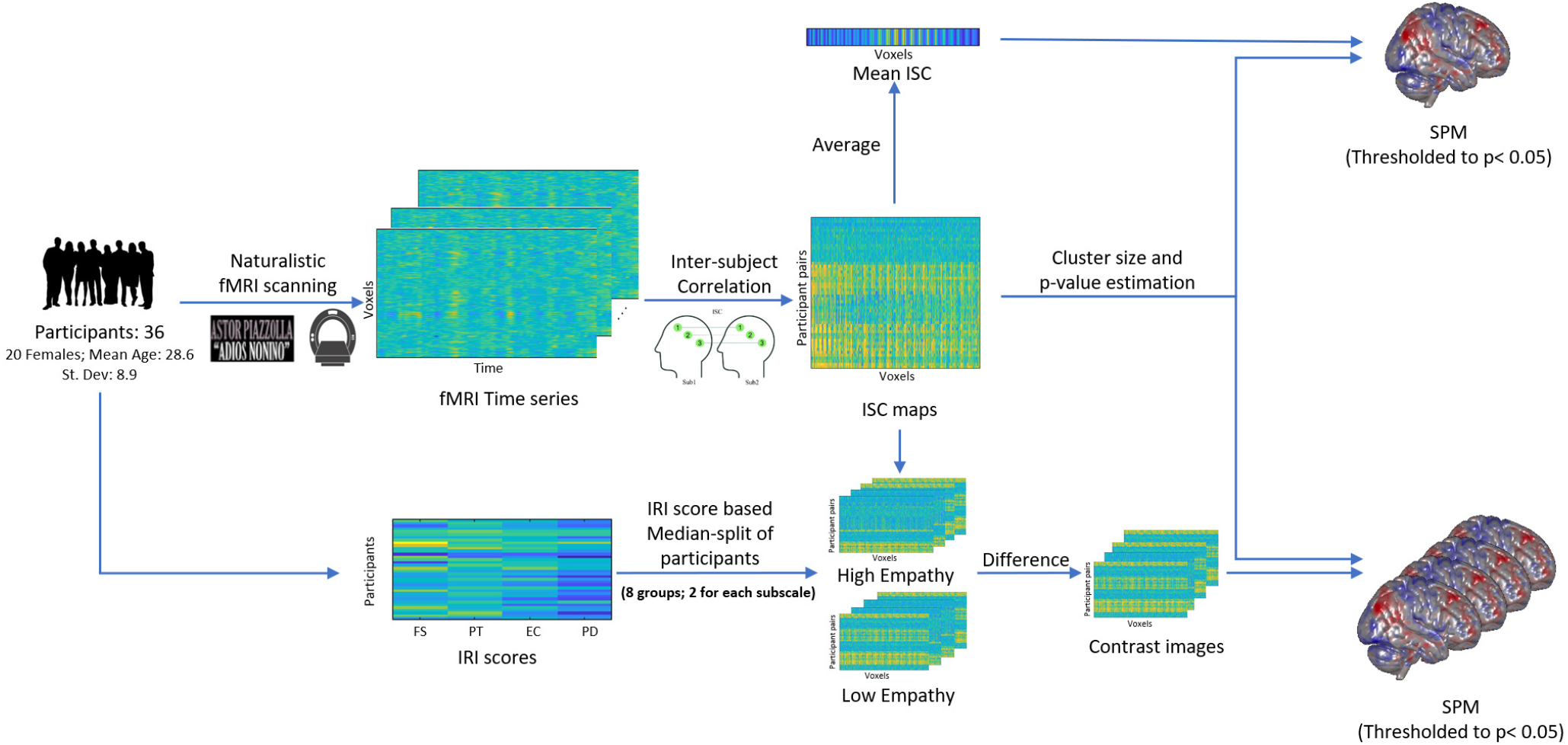
Schematic representation of the Inter-subject correlation analyses.

### Graph based analyses

To understand patterns of functional connectivity from the fMRI time series, we chose to model each participant’s fMRI response as a fully-connected, directed and weighted graph. This allows for the use of various graph-theory based measures to quantify both local and global functional connectivity.

#### The global view: Centrality

Centrality, a widely used graph measure, is a way of quantifying the relative importance of the role played by some nodes in a network (63). Centrality measures have been used in the past to study networks in various fields ranging from social network analysis (64) to studying the spread of epidemics (65). More recently, Centrality has been used as a tool in the network analysis of the human brain (66, 67). Graph centrality is an umbrella term, consisting of various methods such as Node-degree centrality, Betweenness centrality, Eigenvector centrality etc., all of which highlight the important nodes in a network with varying success depending on the network topology (68). The choice of the centrality measure employed would therefore have a significant influence on the results of the study.

One of the more popular methods to measure node centrality - Eigenvector centrality (EVC) is calculated by computing the first eigenvector of a non-negative functional connectivity (similarity) matrix. EVC has seen relative success in the study of social networks (69), since it calculates centrality based not only on the number of neighbouring nodes but also on the centrality of the neighbours unlike other methods such as degree centrality (70). This recursive property allows EVC to reflect global properties within the network. Studies have also shown that the human brain network shows small-world properties very similar to those of social networks (71), making EVC a relevant tool in the study of functional connectivity in the human brain (72). These properties make EVC the appropriate choice for the current study.

In the current study, voxel-wise EVC was computed for each participant, essentially modelling each voxel as a separate graph node. This was done by first generating each participant’s functional connectivity matrix from the fMRI data by computing the pairwise Pearson correlation between each pair of voxel time series. Subsequently, to make the functional connectivity matrix non-negative, each entry was incremented by 1. Finally, a power-iteration method (73) was used to compute the first eigenvector for each participant’s functional connectivity matrix (Refer to Eq. 1). The result was a single EVC brain map for each participant.

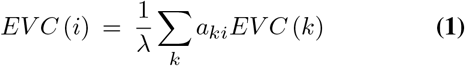

Where:

*EV C*(*i*) = Eigenvector centrality value of the *i*th node

*k* = Denotes all the neighbours of the *i*th node

*a*_*ki*_ = Edge weight (similarity matrix value) between nodes *k* and *i*

*λ* = Constant factor (Eigenvalue)

The resultant EVC maps were then correlated voxelwise with the participants’ IRI scores, using Spearman’s partial correlation due to the presence of significant correlations amongst the various IRI subscales. The correlation maps were thresholded to p<0.05 to retain only the voxels having statistically significant correlation between centrality and IRI scores. To correct for multiple comparisons, we used cluster size thresholding wherein the respective thresholds were obtained from a null distribution obtained via a permutation test. Specifically, we performed 1000 iterations, in which the IRI scores were randomised (with replacement) followed by correlation with EVC values and recording the observed cluster sizes of significant correlations. Cluster sizes were calculated based on the resulting null distribution. An overview of the analysis can be seen in Figure 2.

**Fig. 2.**
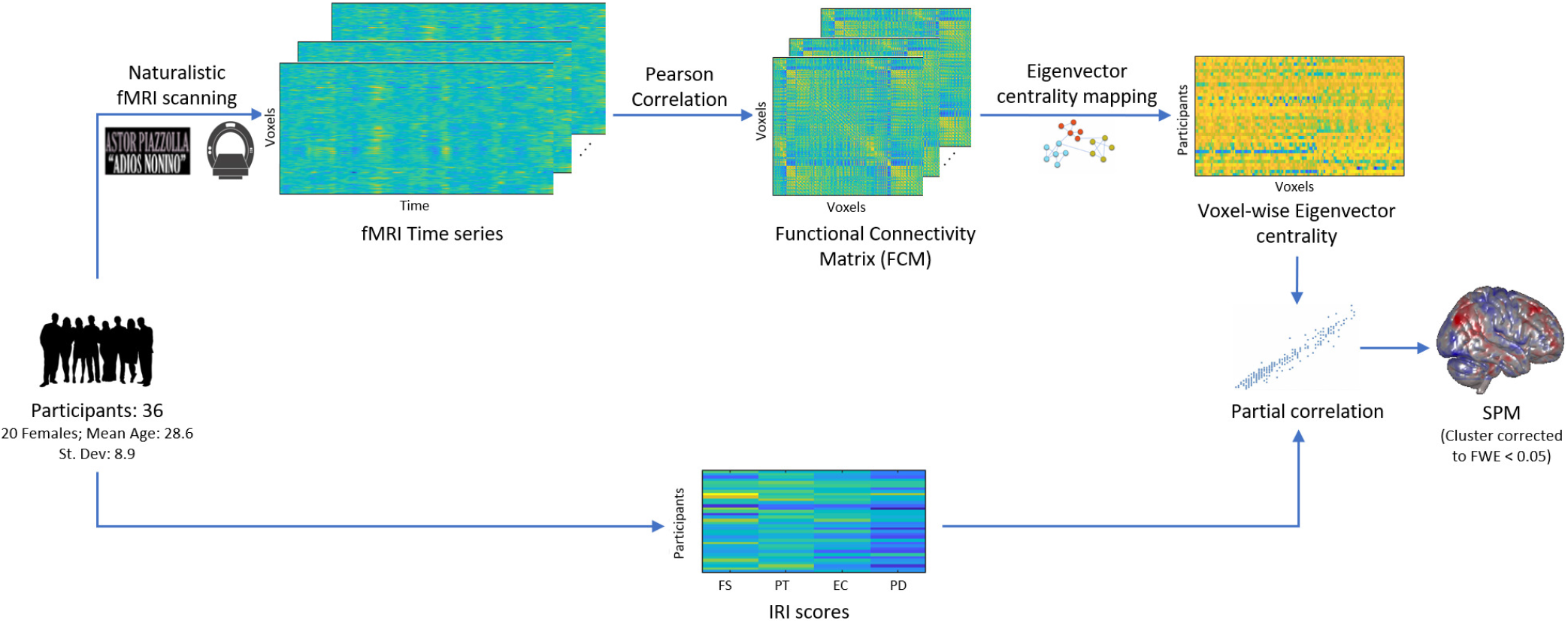
Schematic representation of the Eigenvector centrality analysis.

#### The local view: Modularity

As an exploratory analysis to understand localised connectivity patterns, we employed a graph-based technique to quantify the modularity of neuroanatomically-defined regions of the human brain. Modularity in a graph is a structural measure that quantifies how strongly the network is organised into various modules or subnetworks. In short, a module exhibits higher modularity if the strength of connections within the module is higher than the connections to nodes outside the module. To date modularity in fMRI has only been explored through community detection algorithms wherein one seeks out a modular division of a set of regions into semi-independent communities (74, 75). This is generally accomplished by modelling each atlas-based region as a single node in a graph. Although this approach allows for the identification of a modular subdivision of the various brain regions, it fails to account for the modular organisation within these anatomically defined atlas regions. Moreover, the low resolution (in the order of 100s of nodes) afforded by this approach limits our understanding of the underlying connectivity patterns. Our approach aims to overcome these limitations by both increasing the resolution as well as accounting for the intra-regional connectivity using weighted and signed edges, thereby providing us with a measure for localised connectivity.

In the current study, the results of the EVC analysis were used to select ROIs. This was done by identifying the AAL atlas region which had maximal overlap with each cluster from our EVC analysis. Each of these atlas-based ROIs were then considered as a separate module, whose modularity values were then computed. To reduce the computational burden of constructing the full resolution graph for the measurement of modularity, each participant’s fMRI data was spatially downsampled from 2 × 2 × 2 mm3 sized voxels to 4 × 4 × 4 mm3, resulting in a total of 28,452 voxels. Thereafter, a weighted, signed graph was modelled from the downsampled fMRI time series for each participant, with each voxel as a graph node. The functional connectivity matrix (FCM) was then generated for each participant by computing the pairwise Pearson’s correlation between every pair of nodes. The FCM was subsequently used as the weights of the edges between every pair of voxels, with the sign of the weight denoting the direction of the edge.

#### Calculating Modularity

An overview of the analysis is presented in Figure 3. The various ROIs selected as modules have been outlined in Table 6. With the graph constructed and the modules defined, we proceeded to calculate the Region-wise modularity score of each of the modules defined above. The Modularity measure employed by us was based on the method outlined by Gomez et. al (76), which is a signed and weighted extension of the Newman Modularity measure (77). We chose the Gomez method due to its capability in differentiating between positive and negative edges in the calculation of modularity, allowing us to examine the network with more detail. In the context of human brain connectivity, a positive edge between two voxels denotes that an increase in activation in one of the voxels is accompanied by a proportional increase in in the other whereas a negative edge would imply that an increase in activation in one voxel is accompanied by a proportional decrease in the other. Modularity for a module i is defined by Equation 2a.

**Fig. 3.**
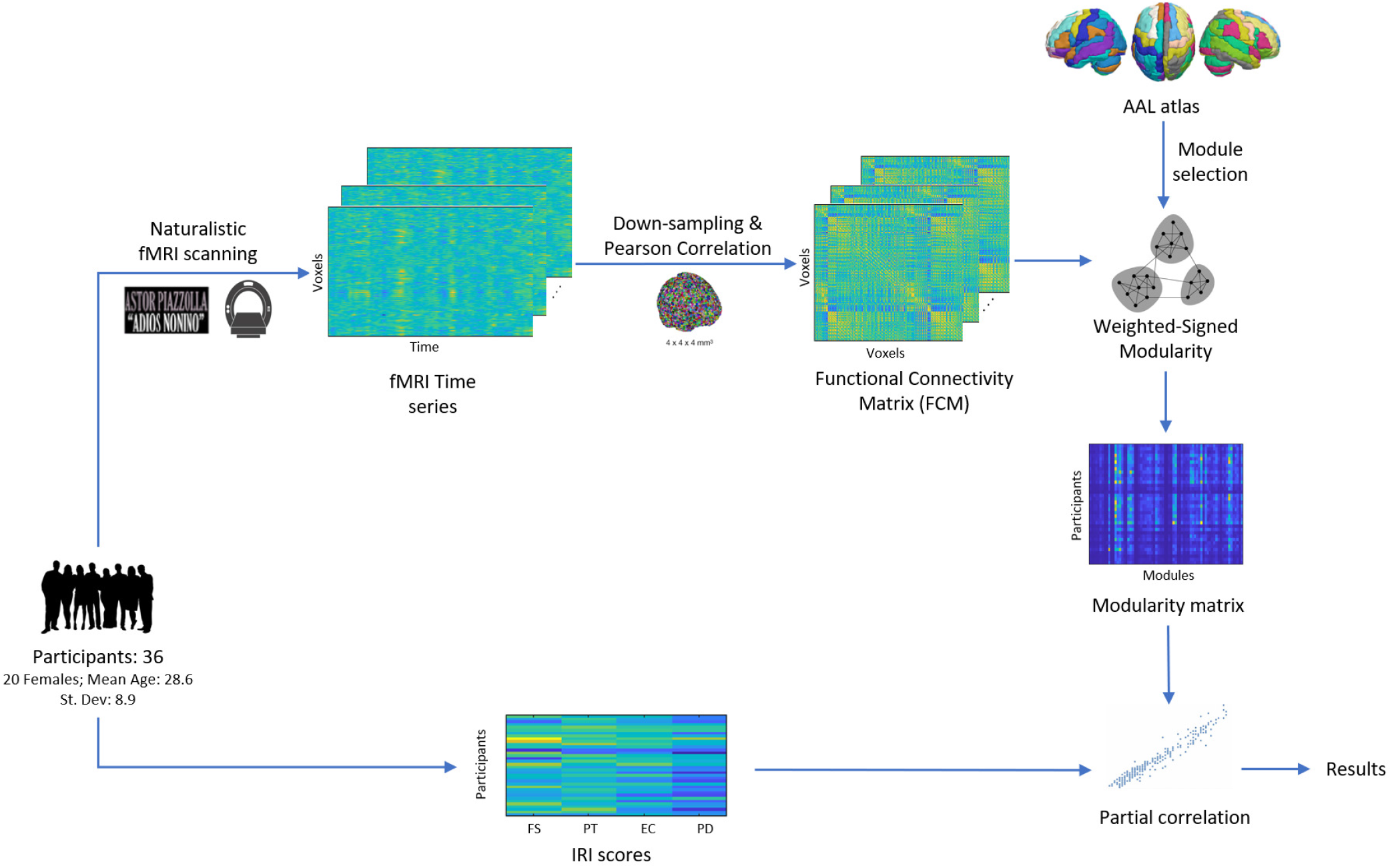
Schematic representation of the exploratory modularity analysis.

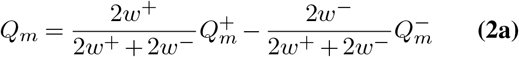

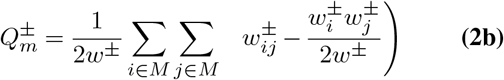

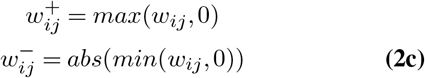

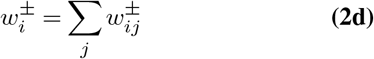

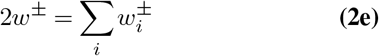

Where:

*m* = Modules in the network

*Q*_*m*_ = Weighted modularity of the *m*th module

*i, j* = Nodes (Downsampled voxels) of the graph

*M* = Set of nodes making up the *m*th Module

*w*_*ij*_ = Weight/Strength of the edge between nodes *i* and *j*

*w*_*i*_, *w*_*j*_ = Node Strength of the *i*th and *j*th nodes; *w*_*i*_ = ∑_*j*_ *w*_*ij*_

2*w* = Total strength of the graph nodes; 2*w* = ∑_*i*_*w*_*i*_

The result of this was a vector of module-wise modularity values for each participant’s fMRI time series. These values were then correlated with participants’ IRI scores using partial Spearman correlation, the results of which have been reported in the next section.

## Results

### Behavioural Measures

The summary of the participants’ demographics and behavioural ratings have been reported in table 1. Lilliefors test (as well as Jarque-Bera goodness-of-fit test) for normality showed that participants’ IRI scores were normally distributed across all 4 subscales (FS, PT, EC, PD) at a 5% significance level. Moreover, Pearson’s correlation performed between the IRI subscales showed a statistically significant correlation between the subscales, which is as expected based on previous studies (55) (Table 2). Hence, this further justifies our use of partial correlations to elicit the differences between the subscales in functional network organisation.

**Table 1.** Summary of participants’ demographic data and behavioural measures.

| Measure | Mean | Std. Deviation |
| --- | --- | --- |
| *Count: 36* |  |  |
| Age | 28.6 | 8.9 |
| Fantasy seeking (FS) | 26.08 | 5.63 |
| Perspective Taking (PT) | 26.25 | 3.54 |
| Emotional Concern (EC) | 24.28 | 3.84 |
| Personal Distress (PD) | 20.81 | 4.97 |
| IRI Total | 97.42 | 13.76 |
| Liking | 2.38 | 0.87 |
| Familiarity | 3.58 | 1.27 |

**Table 2.** Spearman’s correlation between participants’ scores on the four IRI subcales.

| Spearman's Rho | FS | PT | EC | PD |
| --- | --- | --- | --- | --- |
| FS | 1.00 |  |  |  |
| PT | 0.19 | 1.00 |  |  |
| EC | 0.39* | 0.45** | 1.00 |  |
| PD | 0.34* | 0.48** | 0.82*** | 1.00 |

Mann-Whitney U test performed for the Liking and Familiarity scores as well as years of musical training between low and high empathy participants revealed that the distributions for the two groups did not differ significantly at the 5% significance level.

### Inter-subject correlation

#### Mean ISC

As visible in Figure 4 and Table 3, statistically significant clusters showing high mean ISC were found centered around the bilateral auditory cortices (FDR corrected p<0.05). At a less stringent cluster size threshold (95th percentile), an additional cluster was found in the right cerebellum.

**Table 3.** Peak voxels and correlation values for clusters with significant mean ISC. FDR corrected p<0.05; Cluster-size corrected at FWE 0.01.

| Region | Hemisphere | Cluster size | ISC value | MNI (x,y,z) |
| --- | --- | --- | --- | --- |
| Superior Temporal Gyrus,<br>Heschl's gyrus | Left | 180 | 0.17 | -52, -12, 0 |
| Superior Temporal Gyrus,<br>Heschl's gyrus | Right | 654 | 0.20 | 50, -14, 2 |

**Fig. 4.**
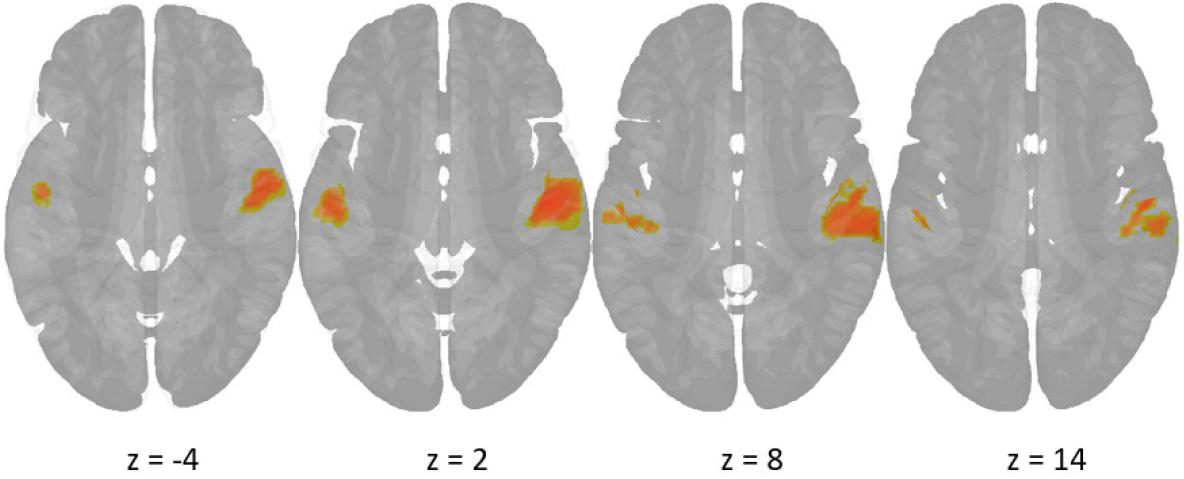
Regions with significantly increased mean ISC scores (p < 0.05). Red - positive ISC.

#### ISC difference between high and low empathy participants

We found significantly greater mean ISC for the high FS group in clusters belonging to the left auditory cortex (middle temporal gyrus) extending to the superior frontal gyrus, sensorimotor regions such as the precentral gyrus, parts of the precuneus and angular gyrus as well as Crus I and II of the cerebellum. In the right hemisphere, we found clusters in parts of calcarine fissure extending to the occipital gyrus, OFC and inferior/superior parietal gyri. (Refer to Figure 5 A, Table 4)

**Table 4.** Peak voxels and ISC difference for significant clusters modulated by IRI scores. FDR corrected p<0.01, Cluster size correlated at FWE 0.05

| Left Hemisphere |  |  |  | Right Hemisphere |  |  |  |
| --- | --- | --- | --- | --- | --- | --- | --- |
| Region | n | MNI (x,y,z) | ISC diff. | Region | n | MNI (x,y,z) | ISC diff. |
| **FS** |  |  |  | **FS** |  |  |  |
| *ISC{high} > ISC{low}* |  |  |  | *ISC{high} > ISC{low}* |  |  |  |
| Middle Frontal gyrus, Precentral gyrus | 547 | -26, 20, 38 | 0.107 | Inferior, Superior Parietal gyrus | 124 | 48, -54, 52 | 0.090 |
| Middle Temporal gyrus | 178 | -58, -6, -22 | 0.131 | Calcarine Fissure, Inferior/Middle Occipital gyrus | 97 | 32, -98, 2 | 0.176 |
| Precuneus (L/R) | 90 | -4, -72, 46 | 0.093 | Superior Frontal gyrus | 46 | 8, 44, 46 | 0.143 |
| Superior Frontal gyrus | 76 | -14, 58, 6 | 0.126 | Gyrus Rectus | 41 | 4, 28, -28 | 0.080 |
| Crus I of Cerebellum | 75 | -34, -72, -28 | 0.153 |  |  |  |  |
| Angular gyrus, Inferior parietal gyrus | 72 | -32, -66, 38 | 0.122 |  |  |  |  |
| Crus I, II of Cerebellum | 52 | -20, -84, -28 | 0.124 |  |  |  |  |
| Superior Frontal gyrus | 43 | -22, 24, 62 | 0.123 |  |  |  |  |
| **EC** |  |  |  | **EC** |  |  |  |
| *ISC{low} > ISC{high}* |  |  |  | *ISC{low} > ISC{high}* |  |  |  |
| Middle/Superior Frontal gyrus | 103 | -20, 42, 24 | -0.126 | Inferior/Superior Frontal gyrus | 97 | 18, 50, -8 | -0.163 |

**Fig. 5.**
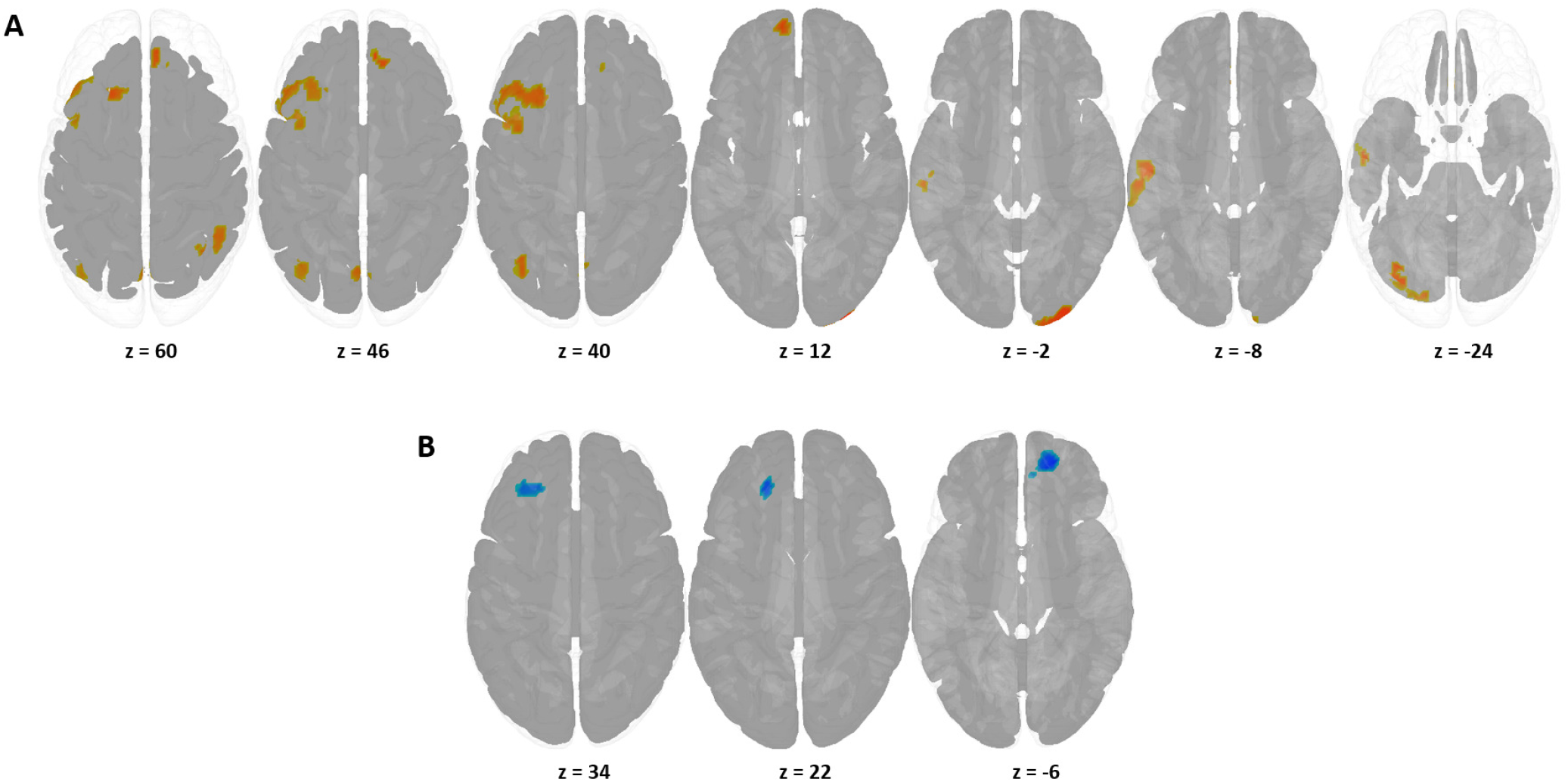
Regions showing significant contrast in ISC between high and low empathy groups for (A) Fantasy Seeking and (B) Empathic concern. Red - ISC for high scoring group > low scoring group, Blue - ISC for low scoring group > high scoring group. Multiple-comparison correction at p < 0.01; Cluster-size corrected at FWE 0.05.

For the EC subscale, the contrast map revealed fewer significant clusters, with greater mean ISC for the low EC groups primarily in and around the bilateral inferior frontal gyrus extending to the superior frontal gyrus (Refer to Figure 5 B, Table 4). We did not find any significant ISC differences for the other subscales.

### Eigenvector centrality

Clusters correlating significantly with empathy scores were found for each of the four IRI subscales. The results are summarised in Table 5.

**Table 5.** Summary of partial Spearman correlation between Eigenvector centrality and IRI scores. p<0.05, Cluster size corrected at FWE 0.05.

| Left Hemisphere |  |  |  | Right Hemisphere |  |  |  |
| --- | --- | --- | --- | --- | --- | --- | --- |
| Region | n | MNI (x,y,z) | z-val | Region | n | MNI (x,y,z) | z-val |
| **FS** |  |  |  | **FS** |  |  |  |
| *Negative* |  |  |  | *Positive* |  |  |  |
| Middle Temporal Gyrus, Inferior temporal gyrus | 193 | -60, -28, -16 | -4.73 | Precentral gyrus, Superior frontal gyrus | 100 | -20, -26, 62 | 4.07 |
| Middle Temporal Gyrus | 158 | -62, -46, 8 | -4.16 | WM tract between MFG and ACC | 68 | 24, 34, 22 | 4.13 |
|  |  |  |  | *Negative* |  |  |  |
|  |  |  |  | Crus I, Crus II of Cerebellum | 290 | 16, -84, -28 | -4.34 |
| **PT** |  |  |  | **PT** |  |  |  |
| *Positive* |  |  |  | *Positive* |  |  |  |
| Posterior Corpus Callosum | 118 | -16, -36, 24 | 3.96 | Supplementary motor area, Superior frontal gyrus, Precentral gyrus, Postcentral gyrus | 237 | 12, -14, 72 | 4.22 |
| Postcentral, Precentral gyrus | 103 | -32, -30, 72 | 4.29 |  |  |  |  |
| *Negative* |  |  |  | *Negative* |  |  |  |
| Putamen, Thalamus and adjoining WM | 45 | -16, -6, 10 | -3.88 | Anterior/Medial cingulate and paracingulate gyrus (R/L), Superior frontal gyrus (R/L) | 381 | 6, 10, 28 | -3.83 |
|  |  |  |  | Caudate nucleus, Globus pallidus, Thalamus | 122 | 10, 2, 0 | -4.19 |
|  |  |  |  | Insula, Inferior frontal gyrus | 112 | 44, 18, -10 | -3.87 |
|  |  |  |  | Thalamus, Brain Stem | 68 | 8, -22, -2 | -4.31 |
| **EC** |  |  |  | **EC** |  |  |  |
| *Positive* |  |  |  | *Positive* |  |  |  |
| Inferior temporal gyrus, Middle temporal gyrus | 86 | -60, -28, -18 | 3.69 | Inferior occipital gyrus, Fusiform gyrus, Crus I of Cerebellum | 181 |  | 4.44 |
| Superior temporal gyrus (Temporal pole), Inferior frontal gyrus | 72 | -42, 18, -16 | 3.58 |  |  |  |  |
| Inferior frontal gyrus, gyrus rectus | 61 | -8, 58, -8 | 3.52 |  |  |  |  |
| *Negative* |  |  |  | *Negative* |  |  |  |
| Precentral gyrus, Paracentral lobule | 164 | -34, -22, 58 | -4.74 | Precuneus and adjacent WM | 315 | 18, -40, 44 | -4.04 |
| Precuneus, Cuneus, Superior parietal gyrus, Superior occipital gyrus | 141 | -14, -58, 32 | -4.49 | Superior occipital gyrus, Precuneus, Angular gyrus | 200 | 26, -66, 38 | -4.54 |
| Fusiform gyrus and WM tract extending to lingual gyrus | 99 | -40, -44, -4 | -4.19 | Middle/Inferior frontal gyrus, Insula, | 119 | 38, 40, 2 | -3.51 |
| Postcentral gyrus, Superior parietal gyrus, Precuneus, Paracentral lobule | 90 | -18, -34, 68 | -3.71 | Insula, Hippocampus | 62 | 38, -12, -10 | -3.56 |
| Inferior frontal gyrus | 52 | -34, 24, 14 | -3.48 | Insula, Inferior frontal gyrus, Putamen | 59 | 40, 10, 6 | -3.49 |
|  |  |  |  | Thalamus | 58 | 24, -24, 6 | -3.26 |
|  |  |  |  | Postcentral gyrus | 51 | 18, -40, 72 | -4.53 |
| **PD** |  |  |  | **PD** |  |  |  |
| *Positive* |  |  |  | *Positive* |  |  |  |
| Inferior frontal gyrus, Insula | 63 | -34, 24, 14 | 3.38 | Middle frontal gyrus, Inferior frontal gyrus | 65 | 38, 42, 2 | 3.25 |

**Table 6.** List of ROIs selected from the AAL atlas for Modularity analysis.

| Selected Modules |  |
| --- | --- |
| Name | Hemisphere |
| Precentral gyrus | L/R |
| Postcentral gyrus | L/R |
| SMA | R |
| Insula | R |
| Thalamus | L/R |
| IFG | L |
| ITG | L |
| MTG | L |
| Precuneus | L/R |
| ACC/MCC | R |
| Cerebellum Crus I | R |
| Cerebellum Crus II | R |
| Gyrus rectus | L/R |
| Temporal Pole | L/R |

High-centrality clusters associated with higher scores on the PT subscale were found bilaterally in sensorimotor areas consisting of the Supplementary Motor Area (SMA), precentral gyrus and the postcentral gyrus Lower PT scores were associated with bilateral clusters around the basal ganglia and thalamus, as well as the right insula and inferior frontal gyrus (IFG). Another cluster belonging to the bilateral CEmN including the anterior and medial cingulate and paracingulate gyri extending to the superior frontal gyrus also show up in participants with low PT scores (Refer to Figure 6 A).

**Fig. 6.**
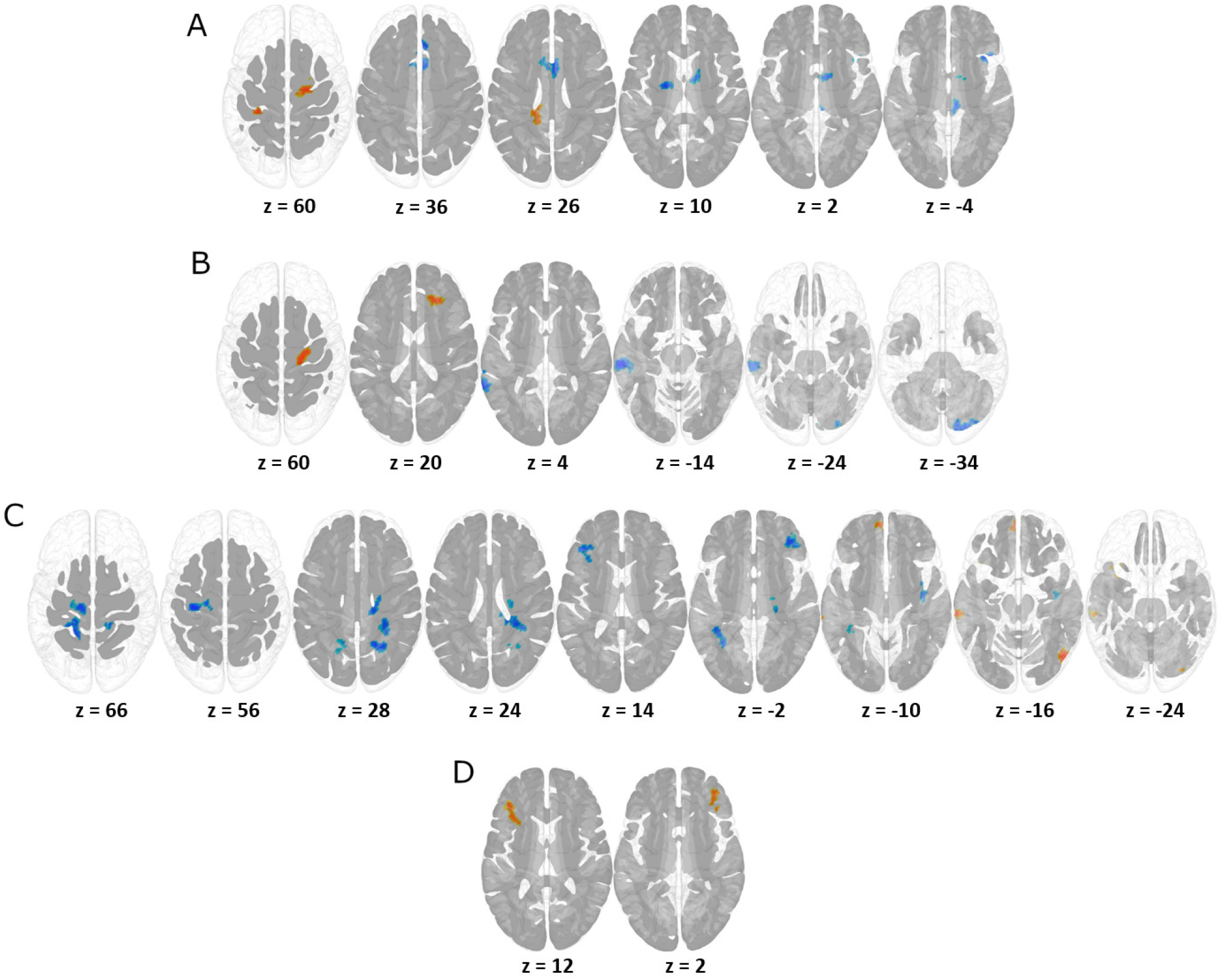
Regions showing significant Spearman partial correlation between voxel-wise eigenvector centrality and empathy scores for (A) Perspective taking (B) Fantasy Seeking (C) Empathic concern (D) Personal Distress. p<0.01, cluster corrected at FWE 0.05. Red - Positive, Blue - Negative correlation.

For the FS subscale, as observed in Figure 6B, high scores were associated with high centrality in the right premotor cortex as well as the white matter tract between the ACC and Middle frontal gyrus. Clusters in left auditory cortex regions (middle/inferior temporal gyri) as well as the right cerebellum (Crus I, II) were found in low FS participants.

The EC subscale showed a higher number of correlated clusters than the other subscales. High scoring participants showed greater centrality in clusters belonging to the left temporal pole (TP) and left orbitofrontal cortex as well as the right occipital lobe extending along the fusiform gyrus to the cerebellum. On the other hand, lower scores were associated with greater centrality in multiple clusters centered bilaterally around the precuneus, primary motor cortex) and IFG ; in the right hemisphere centered around the precentral gyrus extending to the paracentral lobule, the left insula and parts of the occipital lobe. Some clusters were also found to extend into subcortical regions such as the hippocampus, thalamus and basal ganglia (Refer to Figure 6C).

Participants’ PD scores were also observed to be positively correlated with centrality in clusters around the left IFG extending to the insula as well as the right MFG/IFG (Refer to Figure 6 D).

### Modularity

Region-wise modularity scores for all participants ranged from an order of 10^−7^ to 10^−3^, showing a low but highly variable modularity. An overview of the statistically significant partial correlations between modularity and IRI scores can be seen in Table 7 and Figure 7. We found a positive correlation between FS and modularity for the right gyrus rectus (part of OFC). An inverse trend is observed for the PT subscale, which shows a negative correlation with modularity in the right gyrus rectus. Additionally, PT also showed a near-significant negative correlation (p=0.061) with the right SMA and Insula as well as the left IFG, Precuneus and TP.

**Table 7.** Summary of significant partial Spearman correlation (p<0.05) between Modularity and IRI scores.

| Left Hemisphere |  | Right Hemisphere |  |
| --- | --- | --- | --- |
| Module | Spearman's Rho | Module | Spearman's Rho |
| *FS* |  |  |  |
| Gyrus Rectus | 0.35 |  |  |
| *PT* |  |  |  |
| Gyrus Rectus | -0.35 |  |  |
| *EC* |  |  |  |
| Post-Central gyrus | 0.36 | SMA | 0.36 |
| Pre-Central gyrus | 0.39 | Thalamus | 0.44 |
| Thalamus | 0.37 |  |  |
| Temporal Pole | 0.35 |  |  |

**Fig. 7.**
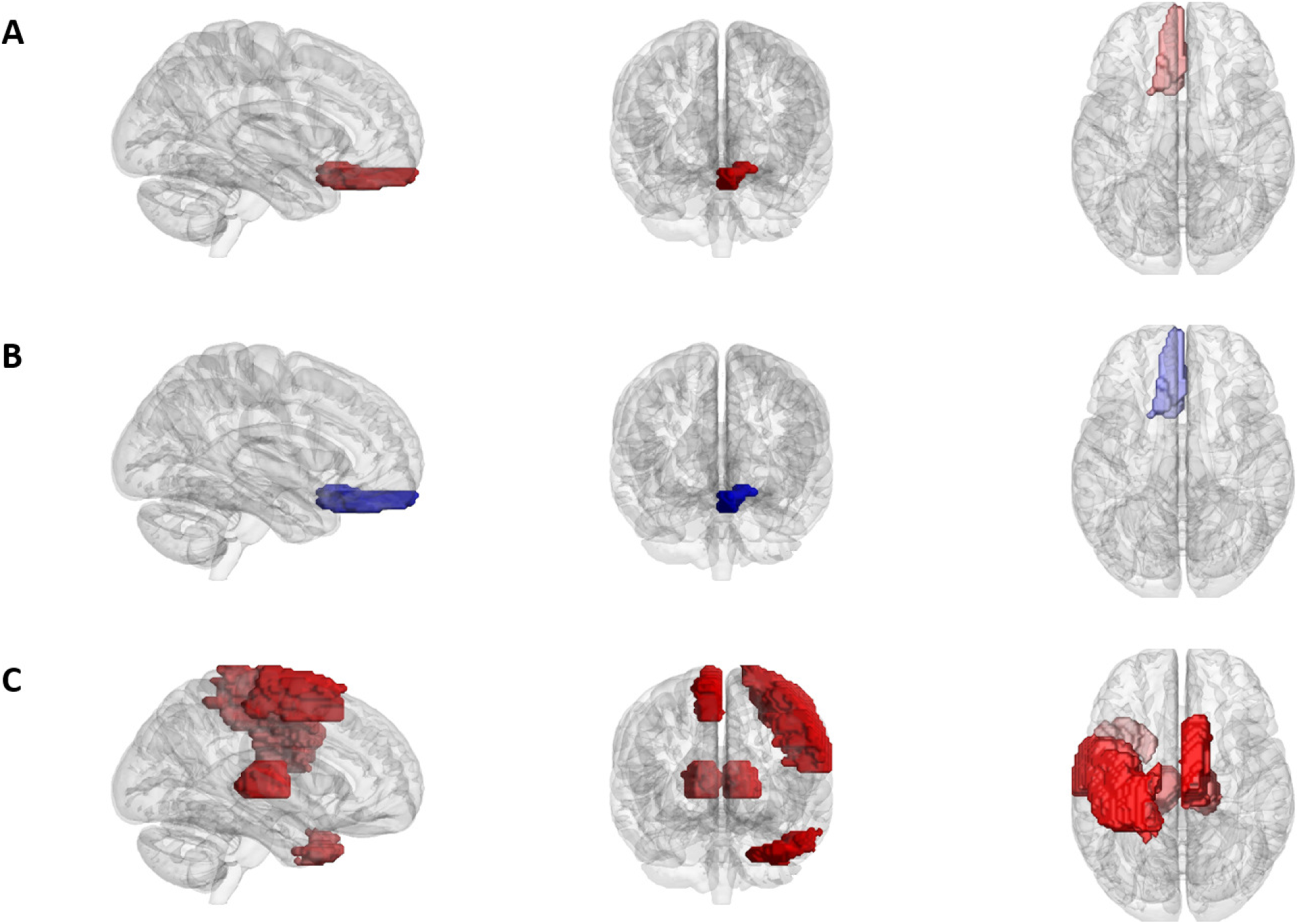
Modules displaying significant correlation between Modularity and Empathy scores for (A) Fantasy Seeking (B) Perspective taking (C) Empathic concern. Red - Positive, Blue - Negative correlation.

Similar to the trend observed in the EVC results, the EC sub-scale was found to be correlated with modularity in a larger number of modules than the other subscales. Positive correlation between modularity and EC were found in the left pre-/post-central gyrus and the right SMA, which belong to the sensorimotor cortex as well as the left TP and bilateral thalamus, which are more integrated with the limbic system. We did not find any significant correlations with modularity scores for the PD subscale.

## Discussion

In the current study, using global and local measures of functional connectivity, we investigated the modulatory effect of trait empathy on whole-brain organization during continuous music listening. Our findings suggest that affective and cognitive empathy differ from each other in the way that they shape whole brain network connectivity, with high cognitive empathy being associated with increased functional centrality in the sensorimotor areas and high affective empathy with increased functional centrality in the OFC and TP, areas involved in the encoding of the affective value of the music, as well as in parts of the occipital lobe and cerebellum. Additionally, we also observed increased global connectivity in the insula and subcortical regions for participants scoring low on both empathy components. Moreover, results from our ISC analysis replicate findings from previous studies such as that of Sachs et al. and further add to the internal consistency and reliability of our data. Furthermore, modularity analyses revealed that EC subscale, that is representative of affective empathy, showed greater intra-regional linkages when compared to FS and PT. Overall, results from our analyses hint at the existence of a left-hemispheric lateralisation in the modulatory effect of empathy, particularly in the OFC and the TP. Our results offer support to previous studies focused on empathy and the processing of music from a naturalistic paradigm and reveal novel findings as discussed below.

### Inter-subject correlation

We used Mean Inter-subject correlation (ISC) to both test the inter-subject consistency of our dataset and to understand how empathy affects ISC at a whole-group level. Our results showed significant mean ISC in voxels belonging to the bilateral auditory cortices (superior temporal gyri, Heschl’s gyri), which is in line with our hypothesis and findings from other studies (39, 47) and is suggestive of the auditory cortex’s role in the affective encoding of the auditory stimuli (Sachs2018). Furthermore, our results also point to lateralisation of this effect, with a greater number of voxels showing high mean ISC in the right hemisphere, which again has been previously observed in other studies using the same paradigm (47).

We subsequently examined the differences in ISC between low- and high empathy participants based on their trait empathy. We found significant differences in mean ISC only for groups divided based on the median scores on FS and EC sub-scales, with the FS subscale showing a more decentralised set of contrasting clusters, in line with the work done by Sachs et al. (39). We observed that participants scoring higher on the FS subscale displayed increased ISC in the left Secondary Auditory cortex, which was also identified by Sachs et al. thereby making this a robust result.

The high FS group also showed significantly greater mean ISC in regions such as the precuneus, parts of the SFG, angular gyrus and the parietal gyrus, all of which belong to the Default Mode Network (DMN), a task negative network that tends to be active in the absence of externally focused attention (78). The DMN has previously been associated with increased activity during day-dreaming and mind-wandering (79); this in addition to the observed cluster in the occipital regions involved in the processing of visual imagery suggest that high FS participants appear to indulge in visual mind-wandering while listening to the current music piece. The music piece used in the current study, Adios Nonino by Piazzolla, is characterised by a melancholic motif in minor mode (80), a musical feature consistently associated with sadness induction (81), supporting the link established between the FS subscale and the enjoyment of sad music in previous studies (18, 39). Additionally, we found high mean ISC for individuals scoring high in FS in the premotor cortex and the cognitive regions of the Cerebellum (Crus I, II) (82). These regions form a part of the cerebello-thalamo-striato-cortical motor pathway, which has previously been shown to be involved in processing of high cognitive load in auditory tasks (83) as well as in the processing of musical rhythm (46).

Contrary to our hypothesis for high EC participants, we failed to find any significant ISC differences in the Thalamus or the Insula. However, we found decreased mean ISC for the high EC participants bilaterally in the frontal regions (IFG/SFG), suggesting that there exists a high degree of variability (in either the duration or intensity) in how high EC participants process musical affect compared to low EC participants.

### Eigenvector Centrality

The eigenvector centrality mapping approach which we used highlights clusters of voxels acting as hubs for communication/coordination, whose centrality reflects participants’ IRI scores. Our results show statistically significant correlations between IRI scores and centrality values in sensorimotor and limbic regions of the brain, which suggests that functional connectivity in empathy networks differs between high and low scoring participants during music listening. We found contrasting trends in centrality between the cognitive and affective scales of the IRI. Overall, in line with our hypothesis, we found that the sensorimotor cortex was associated with increased centrality in participants with higher cognitive empathy and in participants with lower emotional empathy. An unexpected result was the increased centrality in the subcortical regions and Insula in participants with low scores on both affective and cognitive empathy.

We observed a contrasting trend in centrality between EC and FS/PT subscales for clusters belonging to the sensorimotor cortex, specifically the precentral and postcentral gyri and the SMA, with high centrality clusters in this region found in participants scoring high in the FS and PT subscales or scoring low in the EC subscale. This differential role played by the motor areas based in relation to the IRI subscales could be explained by means of motor mimicry playing an important role in cognitive empathy. Scherer and Zentner (21) explained motor mimicry as one mechanism that elicits an emotional response in a listener. This idea of motor mimicry in music is further expanded by Cox’s (84) ‘mimetic hypothesis’, which considers “imitating, covertly or overtly, the observed sound-producing actions of the performers” as key to resonating emotionally with music. It would therefore seem that the sensorimotor cortex plays a central role in people with higher cognitive empathy. More recently, Wallmark et al. (40) also found selective activations in the sensorimotor regions in high FS/PT participants while listening to very short duration sounds, providing support to motor mimicry as an integral part to cognitively empathising with music. The contrasting effect observed with respect to EC in this region could be indicative of a separate neural pathway employed by affective empathy that uses a different centre of coordination as opposed to the sensorimotor cortex. This region also showed greater modularity in high EC participants hinting at more localised processing.

The negative correlation between PT scores and clusters in the subcortical region such as bilateral thalamus and basal ganglia as well the ACC-MCC and right insula indicates that these regions were more central in low scoring participants as they listened to music. The insula and ACC-MCC have been consistently shown to be an integral part of the CEmN across a wide range of activation based studies (37) that predominantly used pain as the stimulus. However, in the context of music listening, Wallmark et al. reported that the ACC was not “a major component in empathy-modulated music processing” (40). Adding to this, our finding that this region is more central in low PT participants seems to suggest that people with low PT are less likely to engage in the Perception-Action coupling that motor mimicry entails, instead relying on the structures in and around the basal ganglia to model their response. A lesion study (85) also found that patients with lesions in the basal ganglia scored significantly lower on the PT scale than the control group. Our results suggest that while listening to music, PT generally recruits a network comprising the basal ganglia and thalamus coupled with the ACC-MCC, but that it is overridden by a more central sensorimotor cortex in the process of motor-mimicry, which is more prevalent in people with high PT. We also observed a contrasting trend in the centrality in clusters around the inferior temporal gyrus, with participants scoring higher on EC showing increased centrality whereas those that scored higher on FS showed reduced centrality. Previous studies have identified this region to perform various functions including higher-order visual processing in addition to processing of auditory irregularities that lead to affective responses (45, 86). Our results therefore seem to suggest that this region may play a central role in the affective processing of music. Additionally, FS also revealed a high centrality in the right cerebellum (Crus I, Crus II) in low scoring participants. Activation in Crus I and Crus II of the right hemisphere has been associated with language processing (82) involving verbal fluency and word generation. One possible interpretation could be that low FS participants processed the music stimuli in terms of language describing the music instead of the affective component in the music.

We find a greater number of central clusters associated with EC than any of the other scales, this may be indicative of a more decentralised network comprising a larger number of hubs associated with the processing of affective empathy. The high centrality clusters found in the left paralimbic region consisting of the orbitofrontal cortex (OFC) and the temporal pole (TP) in high EC participants is indicative of the central role of the region in encoding the affective value of the music. The high centrality observed in our study is in line with previous studies that identify the OFC-TP complex as an association cortex (87) due to its connectivity with the insula, ACC, limbic regions such as the amygdala and hypothalamus (88). Additionally, the OFC-TP complex has also been implicated in the inference of others’ emotional states (89, 90). It is therefore understandable that there exist more well developed central processing hubs in the OFC-TP complex in high EC participants. The observed lateralization in these clusters is also of interest, suggesting a left-hemispheric bias in the OFC-TP complex in empathising with music, this is supported by a previous study (91) that showed drastic changes in affective reactions to auditory stimuli in a patient after the removal of the left TP. Other studies have also implicated a left-hemisphere bias in the TP in the emotional reactions to auditory stimuli such as screaming, crying or laughter (87). Another area, the right inferior occipital gyrus extending to the fusiform gyrus also showed a significant positive correlation with the EC subscale. This region has been implicated in visual imagery (92) as well as in the identification of emotional visual cues (93). Additionally, it has been proposed that empathetic listeners might be more prone to visual imagery, which in turn is reflective of higher susceptibility to musical affect (40).

A number of clusters around the precuneus were found to be negatively correlated with EC scores. The precuneus has previously been identified to constitute the functional core of the Default mode network (DMN) (94), which is a task-negative network (78) that tends to show activity in the absence of any attention-seeking stimuli. The presence of high centrality clusters in this region for people with low EC could be interpreted as lower engagement with the presented stimuli in these participants. Moreover, the precuneus has been linked with the idea of self-consciousness and has shown activations in tasks related to the judgement of one’s own personality traits as opposed to those of others (95, 96). Our finding provides further support and extends the role of the precuneus to perceived emotional states in the self while listening to music. The insula and subcortical regions such as the Thalamus and Basal Ganglia showed a characteristic negative correlation between centrality and both components of empathy. The insula has been shown to be integral in affect processing (97) as well as the processing and integration of interoceptive information with sensory stimuli (98). Additionally, due to its importance in the CEmN, it has also been proposed that the insula may play a role “in the immediate and automatic responses to emotions observed in others, given that it responds to stimuli depicting others in pain regardless of conscious attention or cognitive demand” (39). Studies have also shown that the insula works in tandem with the basal ganglia and thalamus in the bottom-up encoding of affects and moderates the communication with higher-level processing centres such as the sensorimotor cortex and the frontal regions (99). Therefore, the negative correlation observed in these areas for both types of empathy in our study suggests that these regions may work together during music listening as a “default”-empathy coordination center, that moderates communications with more specialised higher-order regions (the sensorimotor cortex for PT or the OFC-TP for EC) and automatically relinquishes its central role to them depending on the stimulus and individual differences.

PD subscale showed positive correlations bilaterally in parts of the IFG associated with recognition of negative valence emotions such as fear, anger or disgust (100, 101). The increased centrality in this region associated with high PD scores can be interpreted in terms of a more central negative affect identification site in these participants, helping them actively be on the lookout for any forms of distressing stimuli.

### Modularity

Our exploratory analysis related to localised connectivity of neuroanatomically defined modules revealed certain interesting patterns which warrant a more thorough examination in future studies. The most prominent of these is a left-hemispheric lateralisation as revealed by the significant correlations observed for all 3 IRI subscales with modularity scores (with the exception of the SMA). This seems to suggest that among the regions identified as being central to empathising to music, namely the sensorimotor cortex and the OFC-TP complex, there is a greater tendency for the left hemisphere to functionally be organised into modular networks based on participants’ empathy scores. Although such a lateralisation effect has previously been observed on lesion studies (85), further research is required to understand its implications in the context of music processing. Additionally, in a similar vein to our EVC analysis findings, we identify a greater number of modules showing significant correlations between Modularity and EC scores compared to the other IRI subscales. This indicates that affective empathy plays a greater modulatory role in the overall functional network connectivity (both global and local) than cognitive empathy, marking it as an important factor to examine in future studies. Due to the higher modularity observed in the left sensorimotor areas (pre-/post-central gyri) in high EC participants paired with the reduced centrality, the authors infer that these regions behave as highly localised modules that take part in global coordination to a lesser extent than is to be expected, cementing what we discussed earlier in terms of centrality. On the other hand, the increased modularity in the TP was also accompanied by increased centrality for high EC participants, meaning that the TP has a greater number of both inter- and intra-modular linkages. It could therefore be inferred that the TP may be acting as a key hub for the coordination, consolidation and subsequent propagation of affective information embedded in the music from across the brain. This is in line with previous research that suggests that the human TP acts as a “cortical convergence zone” (102) that integrates information from the auditory, visual, paralimbic and the default mode networks.

Another interesting observation concerning the two cognitive empathy subscales FS and PT is the contrasting directionality of correlation with modularity score for the gyrus rectus (part of the OFC), with FS exhibiting higher modularity in high scoring participants and PT exhibiting higher modularity in low scoring participants. The increased modularity in this region for high FS participants provides further support to our centrality results by suggesting that this region engages in highly specialised and homogeneous processing of enjoyment of sad music. The reduced modularity associated with PT in this region can be attributed to the high degree of heterogeneity of the frontal regions, allowing it to compartmentalise and process multiple facets of the stimuli simultaneously. In addition to this, the cognitive empathy subscales - PT/FS did not show any correlation with modularity scores in the sensorimotor regions (except for a decreased modularity in the left SMA for FS), suggesting that the cognitive form of motor resonance, that is theorised to primarily take place in these regions does not involve a change in the degree of localised processing.

Overall, these results provide support for the conception of empathy as a complex and multi-faceted phenomenon. They further suggest that differences in individuals’ tendencies to empathize in different ways affect how music is processed in the brain, which may in turn contribute to different emotional responses to heard music. Furthermore, these results lend support to the Cognitive-Affective distinction in how empathy affects the way individuals process various stimuli. Further research may build upon these results by exploring how empathy-related differences in neural processing of music relate to differences in subjective experiences of music, embodied responses to music, and real-time musical interactions. The graph-theory based approach employed in the present study offers insights about global and local network organisation in the human brain while allowing for greater interpretability of the results. Furthermore, a multi-modal approach looking into both structural and functional measures would provide us with a better understanding. However, our results provide support for several key models of empathy and music processing, and merit further investigation into the topic.

## ACKNOWLEDGEMENTS

We would like to thank Brigitte Bogert, Benjamin Gold, Marina Kliuchko, Taru Numminen-Kontti, Johanna Norström, Mikko Heimola, Marita Kattelus, and Toni Auranen. Special thanks to Jyrki Mäkelä, the responsible medical doctor for the study.

## COMPETING FINANCIAL INTERESTS

The authors declare that the research was conducted in the absence of any commercial or financial relationships that could be construed as a potential conflict of interest.

## AUTHOR CONTRIBUTIONS

VM performed the analyses and wrote most of the manuscript; EB collected data and contributed to writing; EC contributed to writing; PT contributed to the analyses and methods; VA is the overseeing author and contributed to writing; PV contributed to shaping the manuscript and interpretations.

## FUNDING

This work was financially supported by the Academy of Finland (author PT, project numbers 272250 and 274037) and Finnish Cultural Foundation (author IB). The Center for Music in the Brain is funded by the Danish National Research Foundation (DNRF project number 117).

## Supplementary Note 1: Demographic data for high and low empathy groups on each IRI sub-scale

Participant demographics for high and low empathy groups based on the median split of the Fantasy Seeking (FS) subscale.

Supplementary Note 1: Demographic data for high and low empathy groups on each IRI subscale Participant demographics for high and low empathy groups based on the median split of the Fantasy Seeking (FS) subscale.
| Measure | High Empathy | Low Empathy |
| --- | --- | --- |
| Age | 28.05±6.44 | 29.47±8.9 |
| Fantasy seeking (FS) | 28.89±1.76 | 23.29±2.54 |
| Perspective Taking (PT) | 26.84±5.36 | 25.23±5.96 |
| Emotional Concern (EC) | 25.63±2.92 | 22.76±4.25 |
| Personal Distress (PD) | 22.78±4.86 | 18.58±4.19 |
| Liking | 2.31±1.24 | 2.47±1.32 |
| Familiarity | 3.73±0.93 | 3.41±0.79 |
| Musical training (Yrs) | 7.42±8.43 | 5.7±8.78 |

Participant demographics for high and low empathy groups based on the median split of the Perspective Taking (PT) subscale.

| Measure | High Empathy | Low Empathy |
| --- | --- | --- |
| Age | 28.16±6.51 | 29.27±8.75 |
| Fantasy seeking (FS) | 26.72±3.75 | 25.77±3.37 |
| Perspective Taking (PT) | 30.33±3.81 | 21.83±3.53 |
| Emotional Concern (EC) | 25.66±3.81 | 22.88±3.42 |
| Personal Distress (PD) | 22.33±5.05 | 19.27±4.52 |
| Liking | 2.55±1.38 | 2.22±1.16 |
| Familiarity | 3.77±0.87 | 3.38±0.84 |
| Musical training (Yrs) | 6.88±7.69 | 6.33±9.48 |

Participant demographics for high and low empathy groups based on the median split of the Empathic Concern (EC) subscale.

| Measure | High Empathy | Low Empathy |
| --- | --- | --- |
| Age | 27.15±5.17 | 30.47±9.52 |
| Fantasy seeking (FS) | 27.36±2.94 | 25±3.82 |
| Perspective Taking (PT) | 27.47±5.9 | 24.52±5.03 |
| Emotional Concern (EC) | 27.26±1.72 | 20.94±2.56 |
| Personal Distress (PD) | 23.78±4.15 | 17.47±3.5 |
| Liking | 2.47±1.34 | 2.29±1.21 |
| Familiarity | 3.57±1.01 | 3.58±0.71 |
| Musical training (Yrs) | 8.21±8.31 | 4.82±8.64 |

Participant demographics for high and low empathy groups based on the median split of the Personal Distress (PD) subscale.

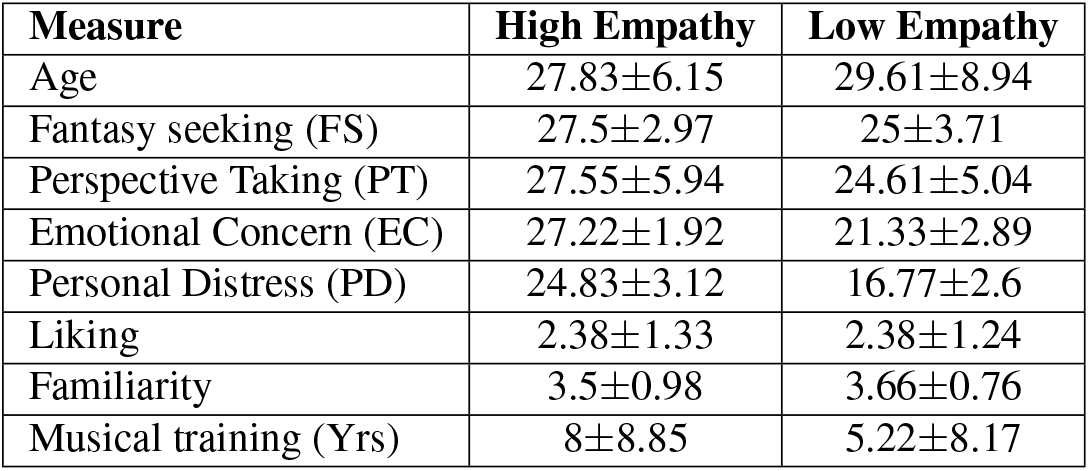

## Notes

### Competing Interest Statement

The authors have declared no competing interest.

